# Matrix-corrected mass spectrometry enables sensitive detection of food allergens and reveals widespread soy contamination of processed foods

**DOI:** 10.1101/231266

**Authors:** Derek Croote, Ido Braslavsky, Stephen R. Quake

## Abstract

The frequent use of precautionary food allergen labeling statements such as “may contain” poses challenges to allergic individuals who rely on such labeling to determine whether a food is safe to consume. We developed a multiplexed liquid chromatography-mass spectrometry assay targeting 14 common allergens in order to survey how frequently these precautionary statements indicate allergen contamination and to assess whether variations in precautionary phrasing affect the likelihood of allergen contamination. A survey of 84 foods revealed how scheduled multiple reaction monitoring (MRM) transition interference derived from complex and heterogeneous sample matrices hinders sensitive analyte detection. As a solution, we developed MAtrix-Dependent Interference Correction (MADIC), an approach to sensitively detect trace peptide quantities through interference identification and stringent peptide quality control criteria. Applying this method, we find frequent contamination of soy in breads and corn flour, and observe additional instances of food contamination with tree nuts, wheat, milk, and egg. In some of these cases, the food had no precautionary labeling for the offending allergen. We also find that only 10% of warning labels are indicative of contamination, and that products with “same facility” precautionary labeling are not necessarily less likely to contain trace amounts of allergens than products labeled “may contain.”

## Introduction

Food allergies are characterized by adverse immunologic reactions to otherwise innocuous food proteins and are estimated to affect 5% of adults and 8% of children (1), with symptoms ranging from vomiting and urticaria to potentially fatal anaphylaxis. Although therapies for desensitization have been developed (2, 3), they do not yet provide a complete cure. Consequently, there is a strong health imperative for reducing the risk of accidental consumption of an allergen by food-allergic individuals. One important step towards this goal is through the labeling of allergen ingredients on commercial food products, mandated by legislation such as the 2004 Food Allergen Labeling and Consumer Protection Act in the U.S or European Directives 2003/89/EC and 2006/142/EC. Numerous countries such as China, Canada, the U.S., as well as the European Union require labeling for products which intentionally contain any of the “big 8” food allergens, namely egg, fish, milk, peanuts, shellfish, soy, tree nuts, and wheat (4, 5). In the Europe Union, this list extends to also include celery, lupin, mustard, sesame, and molluscs. However, these laws do not apply to accidental cross-contamination of allergens into unlabeled foods during manufacturing. While allergen awareness and the implementation of allergen control plans by food manufacturers has increased (6, 7), so has the use of precautionary labeling, such as “may contain” or “produced in a facility that also processes.” The unregulated and frequent use of such statements has confused allergic consumers (8–10) and warrants an assessment of the true underlying risk of food products with precautionary labeling. Such a survey, however, requires an analytical technique embodied by multiplexed allergen detection with sensitivity comparable to eliciting doses in allergic individuals of milligrams or less of allergenic protein in a single serving (11).

Two methodologies exist for the detection of allergenic proteins in food. Antibody-based enzyme-linked immunosorbent assays (ELISAs) are the most common and offer the advantages of high sensitivity and specificity, but suffer when detecting allergens in thermally processed foods (12), differ quantitatively between manufacturers (13), and are difficult to multiplex for testing multiple allergens, although such efforts have been made (14). Liquid chromatography-mass spectrometry is an alternative or complementary analytical technique for allergen detection premised on the tryptic digestion of proteins into peptides that are temporally separated by liquid chromatography prior to interrogation in a mass spectrometer (Fig. 1A). In multiple reaction monitoring (MRM), alternatively known as selected reaction monitoring (SRM), ionized peptides, known as precursor ions, are isolated and fragmented in successive stages of a triple quadrupole mass spectrometer. Resulting product ions are then sequentially isolated in the third quadrupole before striking the detector. Multiple precursor-product ion pairs, known as transitions, are monitored per peptide during a scheduled time window encompassing the peptide’s expected retention time, which maximizes sensitivity by dedicating instrument cycle time to only peptides eluting from the chromatography column at a given time. Peptides are then quantified based on the ratio of native peptide transition areas to transition areas of stable isotope labeled (SIL) peptides spiked into a sample prior to injection. This approach for absolute quantification (15) capitalizes on the identical chromatographic behavior, but different mass, of these internal standards as compared to their native counterparts.

**Fig. 1.**
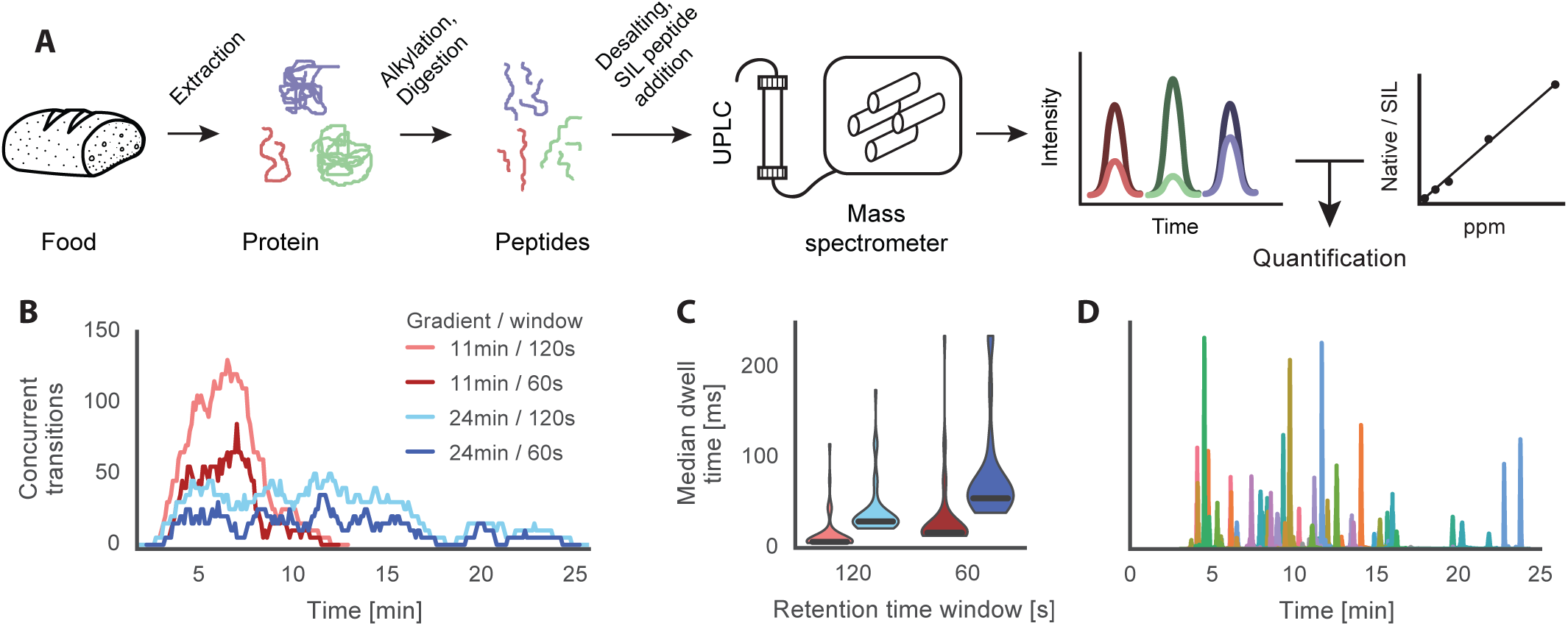
Experimental design and scheduled MRM parameter selection for food allergen detection using mass spectrometry. (A) Protein is extracted from a food sample, alkylated, and digested into peptides, which are then desalted and temporally separated using liquid chromatography prior to interrogation with scheduled MRM on a triple quadrupole mass spectrometer. Quantification is achieved by comparing the ratio of native peptide transition areas over SIL peptide transition areas to a calibration curve. (B) Number of concurrent transitions for 11 min and 24 min elutions with 60 s or 120 s scheduled retention time windows. (C) Violin plots of median dwell times for all 265 transitions partitioned by method configuration in (B). (D) Final 30 min method with a 24 min elution and 60 s scheduled retention time windows for 53 peptides.

One aspect of food allergen detection common to both ELISAs and mass spectrometry is the challenge of sample matrix diversity and complexity; food ingredients each have a unique proteomic profile that, when combined into a food and modified through processing such as baking, equate to innumerable potential matrices. Consequently, although MRM transitions are meant to be specific to the peptide of interest, molecular species in the matrix may unpredictably interfere in transition measurement through co-elution and satisfaction of Q1 and Q3 transmission window resolution constraints (16). This challenge is well-known in the field of clinical proteomics (16, 17), with guidelines for MRM assay development dictating that transitions should be thoroughly screened for matrix-derived interferences (18). This ideal approach is feasible when the diversity of matrices is limited, for example to one or a small number of biological fluids. However, this exclusionary process becomes exceedingly difficult if the assay is to be utilized for many diverse matrices, as is the case with food analyses, where each matrix contains unique interferences. Here we describe a method to address variable interferences derived from multiple sample matrices with minimal loss to overall method sensitivity. We then apply the resulting method to survey commercial foods with and without precautionary labeling for allergen contamination. Our results show widespread contamination of soy in processed food – often without labels – and show that allergen labeling is often not a suitable indicator for estimating allergenicity.

## Results

### Detecting 14 allergens in food using scheduled MRM

We developed a targeted proteomics method on a triple quadrupole mass spectrometer capable of detecting allergenic proteins belonging to the following 14 allergens: almond, brazil nut, cashew, hazelnut, pistachio, walnut, egg, lupin, milk, mustard, peanut, sesame, soy, and wheat. We also assessed the feasibility of targeting gluten proteins derived from barley, oat, rye, and wheat, but found few sufficiently sensitive tryptic peptides, likely due to the absence of alcohol in our extraction solvent that preferentially solubilizes glutens (19–21). For each allergen, we selected two or more tryptic peptides belonging to ideally two or more clinically-recognized allergenic proteins (22) using the Allergen Peptide Browser (23), or via shotgun proteomics of allergen protein extracts (see Materials and Methods as well as Acknowledgments for data availability). Skyline (24) was used to select optimal peptide transitions and establish peptide retention times for synthetic unlabeled peptides and stable isotope labeled (SIL) peptides, which contain ^15^N^13^C-isotopically labeled C-terminal lysine or arginine. To maximize sensitivity, we implemented retention time scheduling, which restricts MRM acquisition to a narrow time window surrounding a peptide’s expected elution from the liquid chromatography column. We used a principled approach to select the scheduling window width in addition to the gradient length under the constraints of cost, throughput, robustness, and sufficient peak sampling. Fig. 1B shows how the gradient length and retention time window affect the number of concurrent method transitions, where higher values corresponds to lower transition dwell times according to the formula 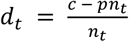, where *c* is the cycle time, *p* is the pause time between transitions, and *n_t_* is the number of transitions at time *t*. We chose a conventional 5 ms pause time and a cycle time of 1.2 s given a 14.4 s median peak width so that at least 10 points per peak were sampled. Compared to an 11 min elution (wash and reequilibration excluded) with 120 s scheduled retention time windows, a 24 min elution with 60 s scheduled retention time windows achieves an eight-fold greater median transition dwell time (Fig. 1C) and was consequently chosen for the final method (Fig. 1D). Through further optimizations, we found that a higher total injection mass improves sensitivity and peak shape (fig. S1A), overnight tryptic digestion is necessary for maximal sensitivity (fig. S1B), eluting with large volumes when desalting maximizes recovery (fig. S1C), and hexane can be used to remove lipids prior to sample protein extraction (25), but may not offer a significant benefit otherwise (fig. S1D-F). For quantification, we used external calibrators with internal standardization by constructing an 8 point calibration curve from 1 to 500 ppm on a food mass basis using a serial dilution of target allergen extracts in a wheat matrix and constant addition of SIL peptides (see Materials and Methods). Allergen detection limits in the wheat matrix, as measured by each allergen’s most sensitive peptide(s), were 5 ppm for 9 allergens, 10 ppm for 1 allergen, and 25 ppm for 3 allergens (table S1). The median R^2^ value of all linear regressions was 0.994. For quantification, the most sensitive peptide “quantifier” was used for each allergen.

### Addressing heterogeneous food matrix interference with MAtrix-Dependent Interference Correction (MADIC)

The complexity of foods combined with their variable composition complicates accurate quantification of allergen contamination. One manifestation of this complexity is interference in MRM transitions, which was observed to be matrix-specific and highly variable in intensity and profile. As depicted in Fig. 2A using three peptides that do not belong to three common food matrices, interference may affect a single transition or multiple, and occur within, crossing, or outside of SIL peptide-defined peak boundaries. These three peptides are representative of the ubiquitous and heterogeneous nature of interference, as illustrated by the extracted ion chromatogram (XIC) for all target transitions (excluding wheat) in an allergen-free wheat matrix (Fig. 2B). Interference is also not limited to a specific instrument at a particular site; interferences were reproduced across sites and instruments (fig. S2) in data from an inter-laboratory plasma protein MRM assay validation study (26).

**Fig. 2.**
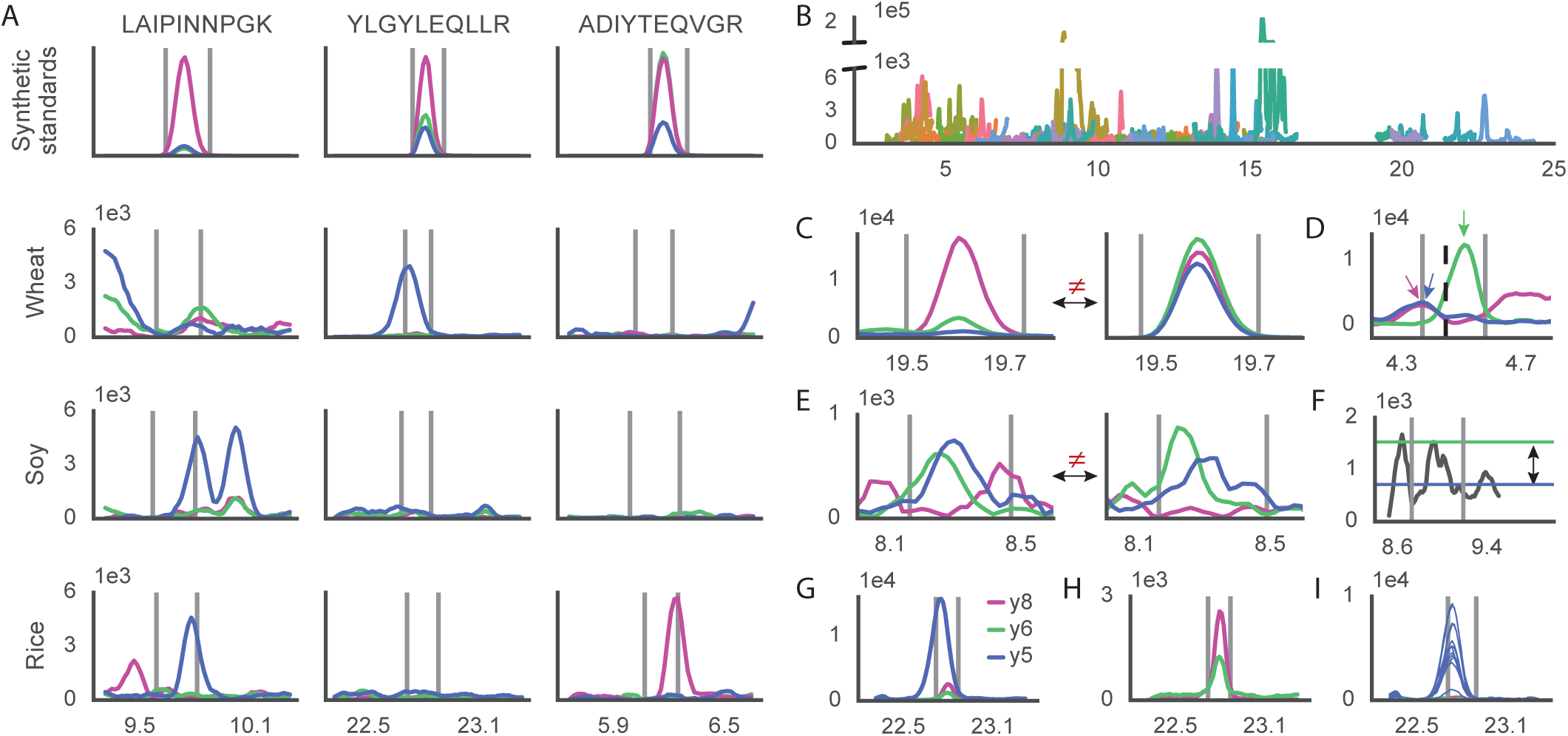
Addressing heterogeneous matrix-specific interference in MRM data. (A) Variable interference in lupin peptide LAIPINNPGK, milk peptide YLGYLEQLLR, and hazelnut peptide ADIYTEQVGR transitions for common food ingredients wheat, soy, and rice. Synthetic standards (upper row) serve as an interference-free reference. Peak boundaries defined by SIL peptides are shown as vertical gray lines. (B) Raw XIC for all non-wheat peptides in a blank wheat matrix. (C-F) Example chromatograms failing quality control criteria. (C) Despite co-eluting within the SIL peptide-defined peak boundaries, transition ratios of the sample (left) disagree with expectations derived from synthetic standards (right). (D) Transitions do not co-elute and individual transition peak retention times (indicated by arrows) do not match the SIL peptide peak retention time (vertical dotted line). (E) Disagreement in transition retention times and transition ratios between duplicate injections. (F) The peak apex (green horizontal line) within the SIL peptide-defined peak boundaries is insufficiently distinguished from the median background (blue line) for the summed transition chromatogram. (G) Interference in the y5 YLGYLEQLLR transition inhibits accurate quantification of 5 ppm milk powder spiked into a wheat matrix. (H) Removal of the y5 wheat interference in (G) rescues peptide confirmation capability. (I) Overlay of twelve bread chromatograms illustrates consistent wheat interference in the y5 YLGYLEQLLR transition.

Trace allergen quantification requires stringent data quality control to address potential false positives arising from heterogeneous matrix interference. We developed a quality control algorithm that enables quantification of allergens from reports and chromatograms exported from Skyline premised on following three quantitative aspects of an interference-free chromatogram: relative transition area ratios similar to synthetic standards, co-elution of all individual target transitions with respective SIL transitions, and a distinguished peak relative to the background. Additionally, the requirement that replicate injections satisfy each of these previous criteria serves as a fourth quality control criterion. Examples of chromatograms identified by our algorithm that diverge from these expected qualities are shown in Fig. 2C-F. This work extends previous efforts limited to transition ratio and replicate filtering (27, 28) by also implementing signal-to-noise and transition retention time matching criteria that together increase confidence in peptide confirmation while also enabling interference identification.

For detecting and quantifying trace amounts of allergens in commercial foods, interference within SIL peptide-defined peak boundaries is especially problematic. For example, compared to a neat synthetic standard (Fig. 2A, center column, first row), the chromatogram of the milk *Bos d 9* (α-s1-casein) peptide YLGYLEQLLR exhibits incorrect transition ratios when attempting to detect 5 ppm milk powder spiked into wheat flour (Fig. 2G). The chromatogram of a blank wheat flour matrix (Fig. 2A, center column, second row) attributes the observed interference in the y5 transition to the wheat matrix, which is corroborated by the fact that this interference is observed in all twelve wheat-based breads tested (Fig. 2I). Removing the y5 transition (Fig. 2H) rescues the remaining y6/y8 transition ratio and therefore could enable quantification of the peptide based on the remaining native to SIL peptide transition area ratio. Importantly, YLGYLEQLLR y5 transition interference is specific to wheat and is not observed in other foods such as soy and rice (Fig. 2A). Consequently, the selective exclusion of high sensitivity transitions with interference in one matrix but not others enables a broadly applicable scheduled MRM method capable of quantifying allergens in a variety of foods, each with a potentially unique interference profile (fig. S3).

### Allergen contamination in commercial foods

We surveyed the precautionary food allergen labeling landscape using scheduled MRM in order to quantify allergen contamination and evaluate if precautionary label phrasing stratified risk. We tested 84 unique food products with a method capable of detecting 14 allergens, with an emphasis on breads, flours, desserts and other snacks (table S2). Some products had no precautionary labeling, while others listed up to six allergens as potential contaminants (Fig. 3A). The value of these labels is unknown and it has been argued that manufacturers use such labeling primarily to avoid litigation (29).

**Fig. 3.**
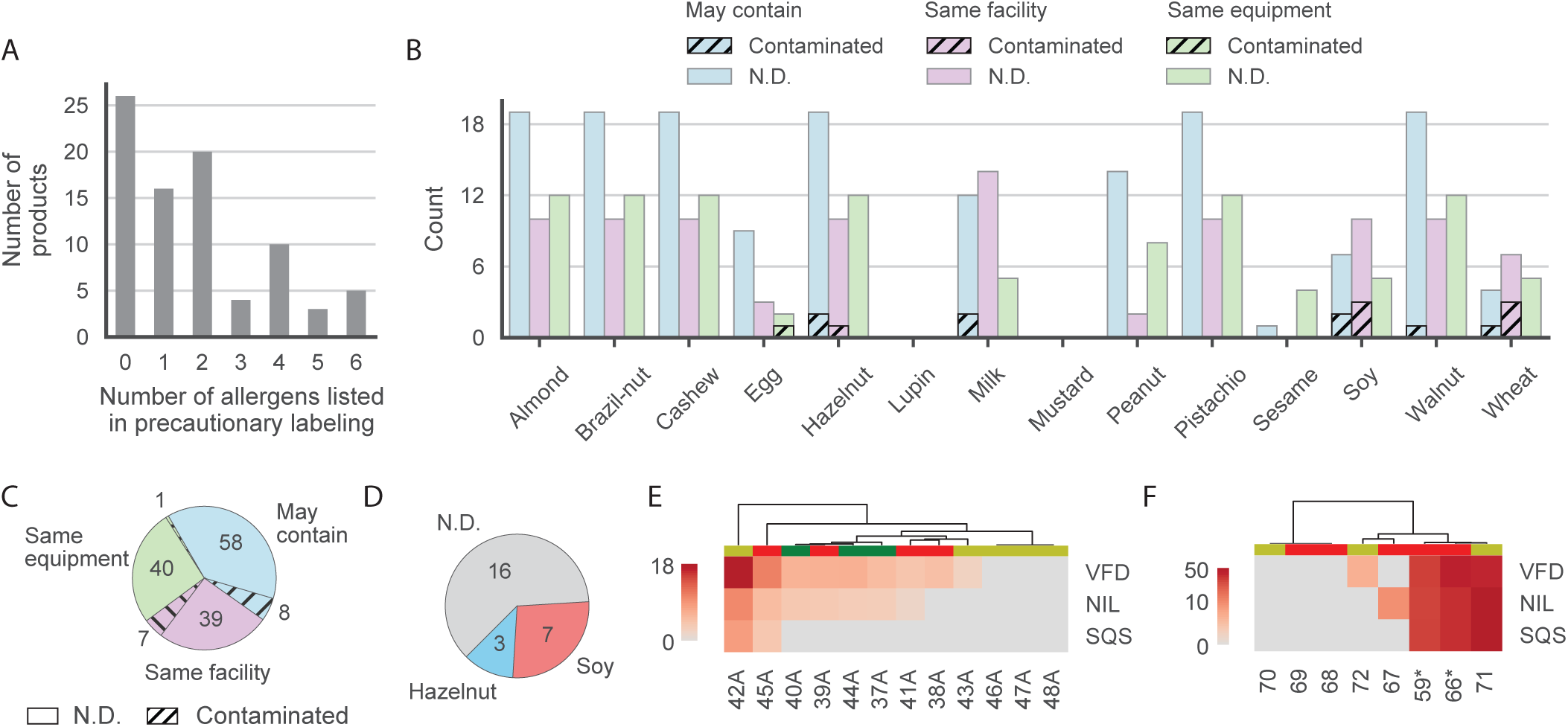
Allergen contamination in commercial foods. (A) Distribution of the number of allergens listed in precautionary labels on 84 commercial food products sampled. (B-C) Contamination in products with precautionary labeling. (B) Instances of allergen contamination, grouped by allergen and precautionary label phrasing type. Label counts for tree nuts have been broadcast to each tree nut. N.D. = not detected. (C) Total number of allergen statements compared to instances of contamination, counting tree nuts as a single allergen. (D) Contamination in products without any precautionary labeling. (E) Heatmap illustrating ppm soy in the crumb (“A” sample number suffix) of twelve bread products. VFDGELQEGR (VFD) is the most sensitive soy peptide and used for quantification, while the others, NILEASYDTK (NIL) and SQSDNFEYVSFK (SQS) are confirmatory. Green column labels indicate soy was listed as an ingredient, while yellow and red column labels indicate presence and absence of soy precautionary labeling, respectively. (F) Heatmap for soy contamination in corn flours. Samples with an asterisk are two separate packages of the same product. Color scale is log_10_ with columns colored as in (E).

We found that precautionary food allergen labeling is overused relative to allergen contamination (Fig. 3B). We examined 84 unique food products, of which 58 had precautionary labeling. Within those labels there were a total of 153 allergens listed; however, we observed only 16 instances of contamination (Fig. 3B-C). In foods with precautionary labeling, we observed the following. Soy, discussed in depth later, is the mostly commonly contaminating allergen, followed by tree nuts and milk. Walnut and hazelnut contaminated 1 and 3 products, respectively, at concentrations of 5 ppm or less and were the only two tree nuts of six found to contaminate any of the products tested, despite similar or better detection limits for the other tree nuts. Milk, which is commonly observed to contaminate chocolate (30–32), was found at 35 ppm [sample 49] and 23 ppm [33] in two snacks containing chocolate chips. The latter, a chocolate chip cookie, elicited an adverse reaction in a milk-allergic individual (private correspondence) and a second package of the same product had similar levels of contamination [38 ppm milk, sample 65]. Interestingly, we found no evidence of milk contamination in a fudge cookie [62] from the same manufacturer, which highlights how allergic consumers can find it difficult to find a trustworthy brand. Wheat was found in one corn flour [71], barley flour [A8], oat flour [A12], and rye flour [A15], while 3 ppm egg was found in one pasta product [55]. Lupin, a flowering plant in the legume family commonly ground into a gluten-free alternative to wheat flour, was not found in any sample, which is consistent with its infrequent use in the U.S. Within the products sampled, peanut was the second-most commonly included allergen in precautionary labeling, and, despite being detected in all products that listed peanut as an ingredient, was not found in any other products. While this is unexpected and may relate to our detection limit or use of a roasted peanut flour as a calibrator (table S1), the trend agrees with previous studies that report lower frequencies of peanut contamination in products with precautionary labeling compared to other allergens such as milk (30, 33).

Precautionary phrasing had numerous slight variations, but could be grouped into the following three general categories: “may contain,” “same facility,” or “same equipment.” Despite how “may contain” was used more frequently than either “same facility” or “same equipment” in the products we tested, the number of contaminated products was similar between “same facility” and “may contain” (Fig. 3C). This result agrees with two previous reports finding no difference in risk between these types of precautionary phrasing for peanut (33) and hazelnut (34). In the context of consumer behavior this is especially worrisome as “same facility” statements are more frequently ignored by allergic individuals and their caregivers when compared with “may contain” statements (8–10).

Allergen contamination is not limited to foods with precautionary labeling: we found soy and tree nuts in products without such labeling (Fig. 3D). Soy was found in two gluten-free pastas [53, 56] at less than 5 ppm as well as in bread and corn flours (discussed below). Hazelnut was found in two cookies [16, 20] and a basil pesto sauce [17] at less than 5 ppm.

### Soy contamination is common in bread and corn flours

We found 20 ppm or less of soy in the crumb (inner portion) of two breads with precautionary labeling, four breads without precautionary labeling, and all three breads labeled as containing soy (Fig. 3E). Soy presence was confirmed in a second set of bread replicates (data not shown). In breads containing soy as an ingredient, soy was present in the form of lecithin, an emulsifier commonly the sole soy-contributing ingredient in large numbers of foods (35). Soybean oil was also present in these samples; however, highly refined oils are exempt from allergen labeling requirements due to their negligible protein content and correspondingly low likelihood of eliciting an allergic reaction (36–38).

We also found a relatively high prevalence of soy contamination in corn flour (Fig. 3F). Of seven unique corn flours tested, four were found to contain soy and of these, only two contained precautionary labeling. Two were found to contain over 40 ppm soy and would consequently elicit an allergic reaction in a greater proportion of allergic individuals than any of the breads examined. For one of these products, we bought a second package and found nearly identical levels of soy contamination (Fig. 3F, samples with asterisks).

### Foods containing soy lecithin are not safe for soy-allergic individuals

Despite the ubiquity of the emulsifier soy lecithin in commercial foods, its allergenicity has been controversial (37, 39). We found soy in 1 of 3 soy lecithin powders (see Materials and Methods), as well as in 24 of 30 unique products containing soy lecithin but no other soy-contributing ingredients (excluding soybean oil). Six samples had no evidence of soy, while 4 products had over 100 ppm (fig. S4). As mentioned previously, the soy in these samples should not originate from soybean oil, but it also might not originate from lecithin and instead could have been introduced during manufacturing or by another ingredient given the growing difficulty of allergen management along increasingly complex and global supply chains. Thus, while the clinical perspective that soy lecithin is minimally allergenic (40) cannot be disproven from these data, our results provide quantitative evidence cautioning against the consumption of lecithin-containing products by soy-allergic individuals due to high levels of soy found in some samples.

### Thermal processing imparts sample-specific allergen losses

For each bread sample, we analyzed both the inner bread portion, known as the crumb, as well as the outer crust of the bread to better understand protein and allergen losses within the context of food processing. The fact that each pair of samples originated from the same dough enables differences to be attributed to temperature- and moisture-dependent effects of baking; for example, the Maillard reaction, which helps impart a brown color to the crust through the reaction of reducing sugars with free amino groups at elevated temperatures (41). These thermal effects significantly reduced the amount of protein extracted from the crust compared to the crumb as measured by microplate colorimetric assay (Fig. 4A). We found soy *Gly m 6* peptide VFDGELQEGR concentration in crust relative to crumb was decreased in a sample-dependent manner even after samples were normalized the same total input injection mass (Fig. 4B). All three wheat allergen peptides depicted a similar trend (Fig. 4C) and losses for the most sensitive wheat peptide TLPTMCR correlated well with those of soy VFDGELQEGR (Fig. 4D). The magnitude of peptide losses varied widely by sample from less than 20% to 100%, which suggests that in bread, losses in the crust are dominated by sample-specific processing effects as opposed to peptide-specific modifications, although it has been suggested that peptides containing lysine may be more susceptible (42).

**Fig. 4.**
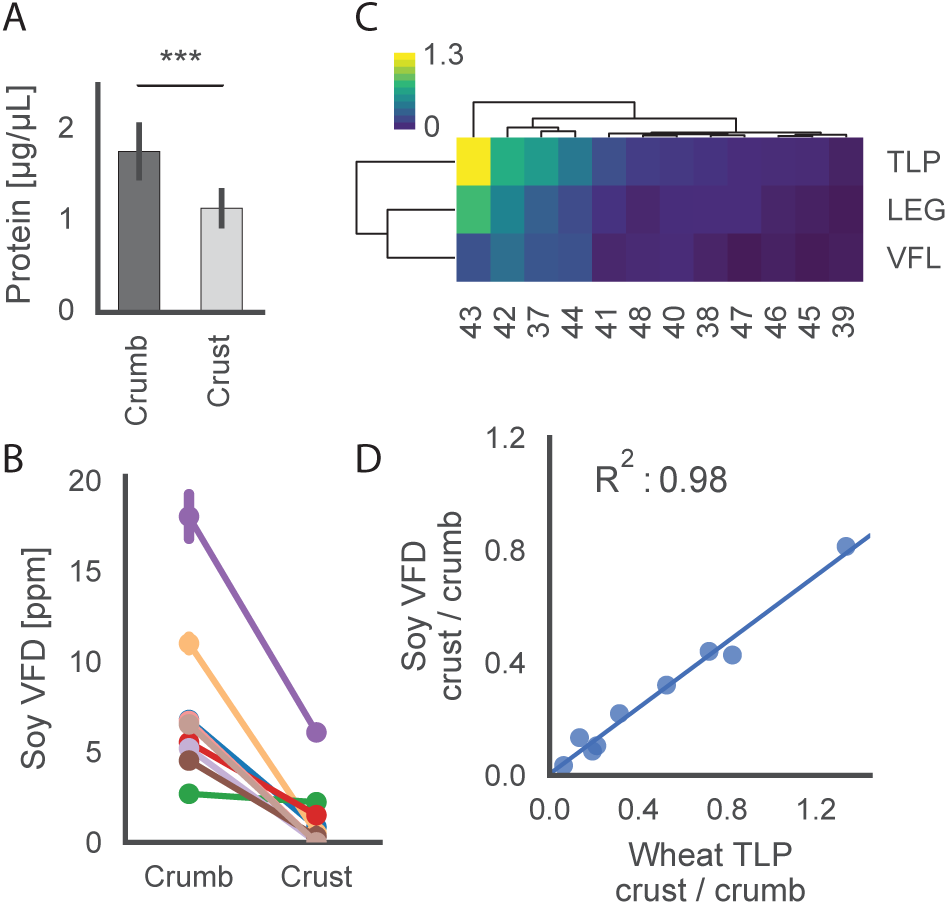
Effects of thermal processing on allergen losses in bread. (A) Protein concentration in paired bread extracts, *** p < 0.001, paired sample t-test, n=12 breads, error bars = sd. (B) ppm soy in paired crust and crumb bread samples as measured by soy *Gly m 6* peptide VFDGELQEGR (VFD). Error bars = sd from duplicate injections, n=9 breads that had detectable levels of soy in the crumb. (C) Heatmap illustrating fraction of SIL-normalized peptide area measured in the crust relative to the crumb for wheat *Tri a 26* peptide LEGSDALSTR (LEG) and *Tri a 36* peptides TLPTMCR (TLP) and VFLQQQCSPVAMPQSLAR (VFL). (D) Crust as a fraction of crumb for soy VFD compared to wheat TLP using SIL-normalized peptide areas.

## Discussion

Parents and allergic individuals will sometimes ignore precautionary statements (8–10, 43) under the assumption that the risk of allergen contamination in these foods is low and phrasing-dependent. We examined the reality of this emergent risk stratification for commercial food products including breads, flours, desserts, snacks and other foods. Such surveys have been performed before in the U.S. (30, 33), U.K. (44), and elsewhere in Europe (34), but have frequently been limited to testing a small number of allergens due to the limited number of commercially available ELISA kits. Our results suggest that while precautionary labeling is rarely indicative of allergen presence, there exists subsets of foods and allergen contaminants which are more likely to co-occur, such as milk in chocolate-containing products and soy in bread and corn flour. Furthermore, we find evidence suggesting that despite consumer behavior, products labeled with “same facility” are not necessarily less likely to contain trace amounts of allergens. In those instances where we do find contamination, the source of contamination is unknown as we have only tested the final output of a complex manufacturing process. Instead, insight into this issue can be gleaned from food recall data gathered by regulatory agencies, such as the U.S. Food and Drug Administration, which point to ingredient mislabeling, cross-contact, improper packaging, improper labeling, omission, and other reasons as common causes of unintended allergen contamination (45). The financial incentive to avoid such recalls (46), combined with efforts to define allergen reference doses (11, 47) and standardize precautionary allergen labeling (48, 49), offers a future path for improved communication of risk to the allergic consumer.

Our computational approach to addressing heterogeneous matrix interference offers advantages over alternative strategies for improving assay selectivity. For example, differential mobility spectrometry (DMS) is an add-on technology to a Sciex QTRAP that utilizes planar electrodes in the mass spectrometer’s source to separate incoming ions based on differences in chemical structure (50). In our hands, the occasional reduction in interference did not outweigh the loss in signal intensity from source hardware differences and longer required pause times between transitions (fig. S5), although we predict this approach may benefit MRM methods with a lower transition density. MRM3 is another alternative to MRM for increasing selectivity whereby the hybrid third quadrupole in a QTRAP instrument, functioning as a linear ion trap, further fragments an MRM product ion into a set of secondary product ions, one or more of which is isolated and detected. This has successfully been applied to a panel of tree nut allergens (51), but the technique cannot be transferred to other triple quadrupole systems and its adoption is further challenged by nonspecific secondary product ions, manual scheduling windows, and long dwell times incompatible with a highly multiplexed method containing SIL peptides. High-resolution mass spectrometers offer an increased ability to discriminate between target analytes and interfering ions of similar mass in complex matrices, but at the cost of dynamic range, sensitivity, and measurement variation compared to MRM on triple quadrupole instruments (52). Parallel reaction monitoring aims to marry the advantages of traditional MRM and high mass accuracy by leveraging instrumentation in which the third quadrupole of a triple quadrupole has been replaced with a high-resolution mass analyzer (53); however, the relative advantages and disadvantages of such an approach compared to existing methods are unknown in the context of multiplexed allergen detection.

For the large majority of allergic individuals, the limits of detection we have demonstrated are sufficiently sensitive to detect reactive allergen quantities. However, depending on the food serving size and allergen, there are about 1-5% of allergic individuals who can react to 0.5 mg or less of allergenic protein (47). The huge dynamic range in allergen sensitivity further illustrates the value of making absolute, quantitative measurements – both for contamination in a given food and in establishing an individual’s limit of sensitivity for an allergen. Approaches such as described here can in principle be used to communicate absolute allergen contamination levels on food labels and consumers would then be able to make choices about whether or not to consume the food based on their own knowledge of their allergen sensitivity, as determined by their doctor.

We developed a method for the detection of fourteen allergens, characterized robust matrix interferences, provided an approach to identify this interference and confidently quantify low abundance peptides, and discovered trends in allergen contamination through a survey of commercial foods that has clear implications for food allergic individuals. The phrasing of labels does not correlate with allergen contamination, and only 10% of allergen statements correspond to detectable amounts of allergen. Conversely, we discovered widespread soy contamination in food without any allergen labels, suggesting a previously unappreciated introduction of soy products into the food supply chain. We envision that after undergoing clinical laboratory standards assay validation with matrix-matched calibration curves constructed using incurred samples, the method we have developed here can be used by consumer organizations, regulatory agencies, or other third parties to verify and catalogue allergen contamination in a broad range of food products.

## Acknowledgments

Shotgun proteomics data have been deposited to the ProteomeXchange Consortium via the PRIDE (54) partner repository with the dataset identifier PXD007688 (http://dx.doi.org/10.6019/PXD007688). Code and examples for matrix-dependent interference correction are available on Github: https://github.com/dcroote/madic.

We would like to acknowledge the staff at the Vincent Coates Foundation Mass Spectrometry Laboratory at Stanford University (SUMS), especially Karolina Krasinska, Allis Chien, and Theresa McLaughlin for helpful discussions related to quantitative method development and instrument performance. This research was supported by the Simons Foundation (SFLIFE #288992 to SRQ), a SUMS seed grant, and the Chan Zuckerberg Biohub. DC is supported by an NSF Graduate Research Fellowship and the Kou-I Yeh Stanford Graduate Fellowship. IB acknowledges support from Stanford University and from The Hebrew University of Jerusalem.

D.C. planned and executed experiments, analyzed the data, developed interference identification and peptide quality control software, prepared figures, and wrote the manuscript. I.B. planned and executed experiments, analyzed the data, and wrote the manuscript. S.R.Q. directed the research and wrote the manuscript.

## References

1. S. H. Sicherer, H. A. Sampson, J Allergy Clin Immunol, in press, doi:10.1016/j.jaci.2013.11.020.

2. W. Yu, D. M. H. Freeland, K. C. Nadeau, Food allergy: immune mechanisms, diagnosis and immunotherapy. Nat Rev Immunol. 16, 751–765 (2016).

3. V. Sampath et al., Deciphering the black box of food allergy mechanisms. Ann Allergy Asthma Immunol. 118, 21–27 (2017).

4. K. J. Allen et al., Precautionary labelling of foods for allergen content: are we ready for a global framework? World Allergy Organiz J. 7, 10 (2014).

5. S. M. Gendel, Comparison of international food allergen labeling regulations. Regul Toxicol Pharmacol. 63, 279–285 (2012).

6. S. M. Gendel, N. Khan, M. Yajnik, A survey of food allergen control practices in the U.S. food industry. J Food Prot. 76, 302–306 (2013).

7. L. S. Jackson et al., Cleaning and other control and validation strategies to prevent allergen cross-contact in food-processing operations. J Food Prot. 71, 445–458 (2008).

8. G. A. Zurzolo, J. J. Koplin, M. L. Mathai, M. K. L. Tang, K. J. Allen, Perceptions of precautionary labelling among parents of children with food allergy and anaphylaxis. Med J Aust. 198, 621–623 (2013).

9. M. Ben-Shoshan et al., Effect of precautionary statements on the purchasing practices of Canadians directly and indirectly affected by food allergies. J Allergy Clin Immunol. 129, 1401–1404 (2012).

10. L. Noimark, J. Gardner, J. O. Warner, Parents’ attitudes when purchasing products for children with nut allergy: a UK perspective. Pediatr Allergy Immunol. 20, 500–504 (2009).

11. K. J. Allen et al., Allergen reference doses for precautionary labeling (VITAL 2.0): clinical implications. J Allergy Clin Immunol. 133, 156–164 (2014).

12. S. Khuda et al., Effect of processing on recovery and variability associated with immunochemical analytical methods for multiple allergens in a single matrix: sugar cookies. J Agric Food Chem. 60, 4195–4203 (2012).

13. S. Jayasena et al., Comparison of six commercial ELISA kits for their specificity and sensitivity in detecting different major peanut allergens. J Agric Food Chem. 63, 1849–1855 (2015).

14. C. Y. Cho, W. Nowatzke, K. Oliver, E. A. E. Garber, Multiplex detection of food allergens and gluten. Anal Bioanal Chem. 407, 4195–4206 (2015).

15. S. A. Gerber, J. Rush, O. Stemman, M. W. Kirschner, S. P. Gygi, Absolute quantification of proteins and phosphoproteins from cell lysates by tandem MS. Proc Natl Acad Sci U S A. 100, 6940–6945 (2003).

16. M. A. Gillette, S. A. Carr, Quantitative analysis of peptides and proteins in biomedicine by targeted mass spectrometry. Nat Methods. 10, 28–34 (2013).

17. S. Gallien, E. Duriez, K. Demeure, B. Domon, Selectivity of LC-MS/MS analysis: implication for proteomics experiments. J Proteomics. 81, 148–158 (2013).

18. S. A. Carr et al., Targeted peptide measurements in biology and medicine: best practices for mass spectrometry-based assay development using a fit-for-purpose approach. Mol Cell Proteomics. 13, 907–917 (2014).

19. M. L. Colgrave, K. Byrne, M. Blundell, C. A. Howitt, Identification of barley-specific peptide markers that persist in processed foods and are capable of detecting barley contamination by LC-MS/MS. J Proteomics. 147, 169–176 (2016).

20. M. L. Colgrave et al., Comparing multiple reaction monitoring and sequential window acquisition of all theoretical mass spectra for the relative quantification of barley gluten in selectively bred barley lines. Anal Chem. 88, 9127–9135 (2016).

21. S. Lock, Gluten Detection and Speciation by Liquid Chromatography Mass Spectrometry (LC-MS/MS). Foods. 3, 13–29 (2013).

22. C. Radauer et al., Update of the WHO/IUIS Allergen Nomenclature Database based on analysis of allergen sequences. Allergy. 69, 413–419 (2014).

23. D. Croote, S. R. Quake, Food allergen detection by mass spectrometry: the role of systems biology. npj Syst Biol Appl. 2, 16022 (2016).

24. B. MacLean et al., Skyline: an open source document editor for creating and analyzing targeted proteomics experiments. Bioinformatics. 26, 966–968 (2010).

25. C. H. Parker et al., Multi-allergen Quantitation and the Impact of Thermal Treatment in Industry-Processed Baked Goods by ELISA and Liquid Chromatography-Tandem Mass Spectrometry. J Agric Food Chem. 63, 10669–10680 (2015).

26. S. E. Abbatiello et al., Large-Scale Interlaboratory Study to Develop, Analytically Validate and Apply Highly Multiplexed, Quantitative Peptide Assays to Measure Cancer-Relevant Proteins in Plasma. Mol Cell Proteomics. 14, 2357–2374 (2015).

27. Y. Bao et al., Detection and correction of interference in SRM analysis. Methods. 61, 299–303 (2013).

28. S. E. Abbatiello, D. R. Mani, H. Keshishian, S. A. Carr, Automated detection of inaccurate and imprecise transitions in peptide quantification by multiple reaction monitoring mass spectrometry. Clin Chem. 56, 291–305 (2010).

29. J. B. Roses, Food Drug Law J, in press.

30. M. P. Crotty, S. L. Taylor, Risks associated with foods having advisory milk labeling. J Allergy Clin Immunol. 125, 935–937 (2010).

31. U.S. Food and Drug Administration, A Survey of Milk in Dark Chocolate Products (2015), (available at https://www.fda.gov/Food/IngredientsPackagingLabeling/FoodAllergens/ucm446646.htm).

32. S. L. Hefle, D. M. Lambrecht, Validated sandwich enzyme-linked immunosorbent assay for casein and its application to retail and milk-allergic complaint foods. J Food Prot. 67, 1933–1938 (2004).

33. S. L. Hefle et al., Consumer attitudes and risks associated with packaged foods having advisory labeling regarding the presence of peanuts. J Allergy Clin Immunol. 120, 171–176 (2007).

34. M. Pele, M. Brohée, E. Anklam, A. J. Van Hengel, Peanut and hazelnut traces in cookies and chocolates: relationship between analytical results and declaration of food allergens on product labels. Food Addit Contam. 24, 1334–1344 (2007).

35. M. M. Pieretti, D. Chung, R. Pacenza, T. Slotkin, S. H. Sicherer, Audit of manufactured products: use of allergen advisory labels and identification of labeling ambiguities. J Allergy Clin Immunol. 124, 337–341 (2009).

36. R. K. Bush, S. L. Taylor, J. A. Nordlee, W. W. Busse, Soybean oil is not allergenic to soybean-sensitive individuals. J Allergy Clin Immunol. 76, 242–245 (1985).

37. Awazuhara, Kawai, Baba, Matsui, Komiyama, Antigenicity of the proteins in soy lecithin and soy oil in soybean allergy. Clin Exp Allergy. 28, 1559–1564 (1998).

38. A. Paschke, K. Zunker, M. Wigotzki, H. Steinhart, Determination of the IgE-binding activity of soy lecithin and refined and non-refined soybean oils. J Chromatogr B Biomed Sci Appl. 756, 249–254 (2001).

39. U. Müller et al., Commercial soybean lecithins: a source of hidden allergens? Zeitschrift für Lebensmitteluntersuchung und-Forschung A. 207, 341–351 (1998).

40. J. D. Kattan, R. R. Cocco, K. M. Järvinen, Pediatr Clin North Am, in press, doi:10.1016/j.pcl.2011.02.005.

41. S. I. F. . Martins, W. M. . Jongen, M. A. J. . van Boekel, A review of Maillard reaction in food and implications to kinetic modelling. Trends Food Sci Technol. 11, 364–373 (2000).

42. G. A. Newsome, P. F. Scholl, Quantification of allergenic bovine milk α(S1)-casein in baked goods using an intact ^15^N-labeled protein internal standard. J Agric Food Chem. 61, 5659–5668 (2013).

43. M. J. Marchisotto et al., Global perceptions of food allergy thresholds in 16 countries. Allergy. 71, 1081–1085 (2016).

44. B. Hirst, “Survey of allergen advisory labelling and allergen content of UK retail pre-packed processed foods” (FS241038 (T07067), Reading Scientific Services LTD, United Kingdom, 2014).

45. S. M. Gendel, J. Zhu, Analysis of U.S. Food and Drug Administration food allergen recalls after implementation of the food allergen labeling and consumer protection act. J Food Prot. 76, 1933–1938 (2013).

46. R. S. Gupta et al., Economic Factors Impacting Food Allergen Management: Perspectives from the Food Industry. J Food Prot. 80, 1719–1725 (2017).

47. S. L. Taylor et al., Establishment of Reference Doses for residues of allergenic foods: report of the VITAL Expert Panel. Food Chem Toxicol. 63, 9–17 (2014).

48. S. Hattersley, R. Ward, A. Baka, R. W. R. Crevel, Advances in the risk management of unintended presence of allergenic foods in manufactured food products--an overview. Food Chem Toxicol. 67, 255–261 (2014).

49. P. J. Turner, A. S. Kemp, D. E. Campbell, Advisory food labels: consumers with allergies need more than “traces” of information. BMJ. 343, d6180 (2011).

50. J. L. Campbell, J. C. Y. Le Blanc, R. G. Kibbey, Differential mobility spectrometry: a valuable technology for analyzing challenging biological samples. Bioanalysis. 7, 853–856 (2015).

51. R. Korte, J. Brockmeyer, MRM(3)-based LC-MS multi-method for the detection and quantification of nut allergens. Anal Bioanal Chem. 408, 7845–7855 (2016).

52. B. Domon, R. Aebersold, Options and considerations when selecting a quantitative proteomics strategy. Nat Biotechnol. 28, 710–721 (2010).

53. A. C. Peterson, J. D. Russell, D. J. Bailey, M. S. Westphall, J. J. Coon, Parallel reaction monitoring for high resolution and high mass accuracy quantitative, targeted proteomics. Mol Cell Proteomics. 11, 1475–1488 (2012).

54. J. A. Vizcaíno et al., 2016 update of the PRIDE database and its related tools. Nucleic Acids Res. 44, D447–56 (2016).

55. R. Korte, S. Lepski, J. Brockmeyer, Comprehensive peptide marker identification for the detection of multiple nut allergens using a non-targeted LC-HRMS multi-method. Anal Bioanal Chem. 408, 3059–3069 (2016).

56. V. Sharma et al., Panorama: a targeted proteomics knowledge base. J Proteome Res. 13, 4205–4210 (2014).

